# Transcription activator WCC recruits deacetylase HDA3 to control transcription dynamics and bursting in Neurospora

**DOI:** 10.1101/2023.02.08.527627

**Authors:** Michael Oehler, Axel C.R. Diernfellner, Michael Brunner

**Affiliations:** Heidelberg University Biochemistry Center, Im Neuenheimer Feld 328, D-60120 Heidelberg, Germany

**Keywords:** histone deacetylase 3, HDAC3, HDA3, transcription activation, transcription repression, transcriptional bursting, White Collar Complex, light-induced transcription

## Abstract

RNA polymerase II initiates transcription either randomly or in bursts. We examined the light-dependent transcriptional activator White Collar Complex (WCC) of *Neurospora* to characterize the transcriptional dynamics of the strong *vivid* (*vvd*) promoter and the weaker *frequency* (*frq*) promoter. We show that WCC is not only an activator but also represses transcription by recruiting histone deacetylase 3 (HDA3). Our data suggest that bursts of *frq* transcription are governed by a long-lived refractory state established and maintained by WCC and HDA3 at the core promoter, whereas transcription of *vvd* is determined by WCC binding dynamics at an upstream activating sequence. Thus, in addition to stochastic binding of transcription factors, transcription factor-mediated repression may also influence transcriptional bursting.

**TEASER:** Balanced interaction of transcription factor with coactivator and corepressors determines transcription dynamics and bursting.

## INTRODUCTION

Regulated transcription is induced by sequence-specific transcription factors (TFs) that cause the remodelling or removal of nucleosomes in the promoters of their target genes and facilitate the assembly of the general transcription machinery and RNA Polymerase II (Pol II) into an active preinitiation complex (PIC) (*1*). The active state of the promoter is competed by the disassembly of the PIC and eventually the (de)modification and repositioning of the nucleosomes. The active promoter state can be very short-lived, leading to transcription bursts interrupted by transcriptionally inactive phases which can be considerably long (*2-10*). TF binding dynamics are rate-limiting for the frequency of transcription initiation in many cases, and there is compelling evidence that the stochasticity of TF binding is a cause of transcriptional bursting (*8, 11-16*).

Analysis of the transcription dynamics of single gene loci in single cells has shown that the size, duration, and frequency of transcription bursts depend on TF abundance and promoter architecture and are modulated by enhancer activity (*12, 13, 17-19*). Transcriptional bursting is a widespread phenomenon (*4, 7, 18*). Typically, transcriptional bursting is mathematically described by a two state - on-off - promoter model. Simple transcription bursts last a few minutes but can be combined to complex bursts with much longer promoter on-times lasting, in some cases even hours; and similarly, promoter off-times vary from minutes to hours (*4, 7*). Statistical analyses revealed that the promoter off-times are not always entirely stochastic in nature (*4, 20*). Rather, in several cases, the promoter off-time is composed of a period immediately following a transcription burst in which the promoter is refractory to restimulation and a subsequent permissive period in which transcriptional reactivation is determined by the stochasticity of TF binding, which is best described by a three-state promoter model (*4, 20, 21*). Mechanisms underlying transcriptional refractoriness remain elusive.

Sophisticated, single-cell-based approaches allow simultaneous monitoring of the dynamics of TF binding and transcription of engineered reporter genes in the living cell (*12, 13*) but are less suited identifying the components governing bursting. Biochemical or sequencing-based analyses of transcriptional dynamics would in principle be suitable for determining factors and mechanisms underlying these processes but this would require sufficiently tight synchronization of transcriptional bursts in a cell population so that the successive steps of a ‘forced’ transcriptional burst can be temporally resolved and analyzed.

The light-responsive White Collar Complex (WCC) is a core component of the circadian clock of *Neurospora crassa* (*22*) and provides a natural optogenetic TF tool that allows for the quantitative analysis of transcription dynamics in the living cell with high resolution (*20, 21*) (Fig.1A). Light enables immediate activation and dimerization of the WCC and the nuclear concentration of WCC is high enough to ensure rapid initiation of transcription, a prerequisite for generating a population-wide, tightly synchronized transcription wave. Studies revealed that WCC-induced transcription from the *frequency* (*frq*) promoter is rapidly limited by transcription-dependent active repression (*21*). The repressed state of the *frq* promoter is exceptionally long-lived (t_1/2_ ∼45 min). During this period the promoter is refractory to restimulation despite efficient recruitment of new WCC activated by a second LP. In steady state (constant light), the refractory behavior of the *frq* promoter must inevitably lead to transcriptional bursts interspersed by long promoter off-times.

**Fig. 1.**
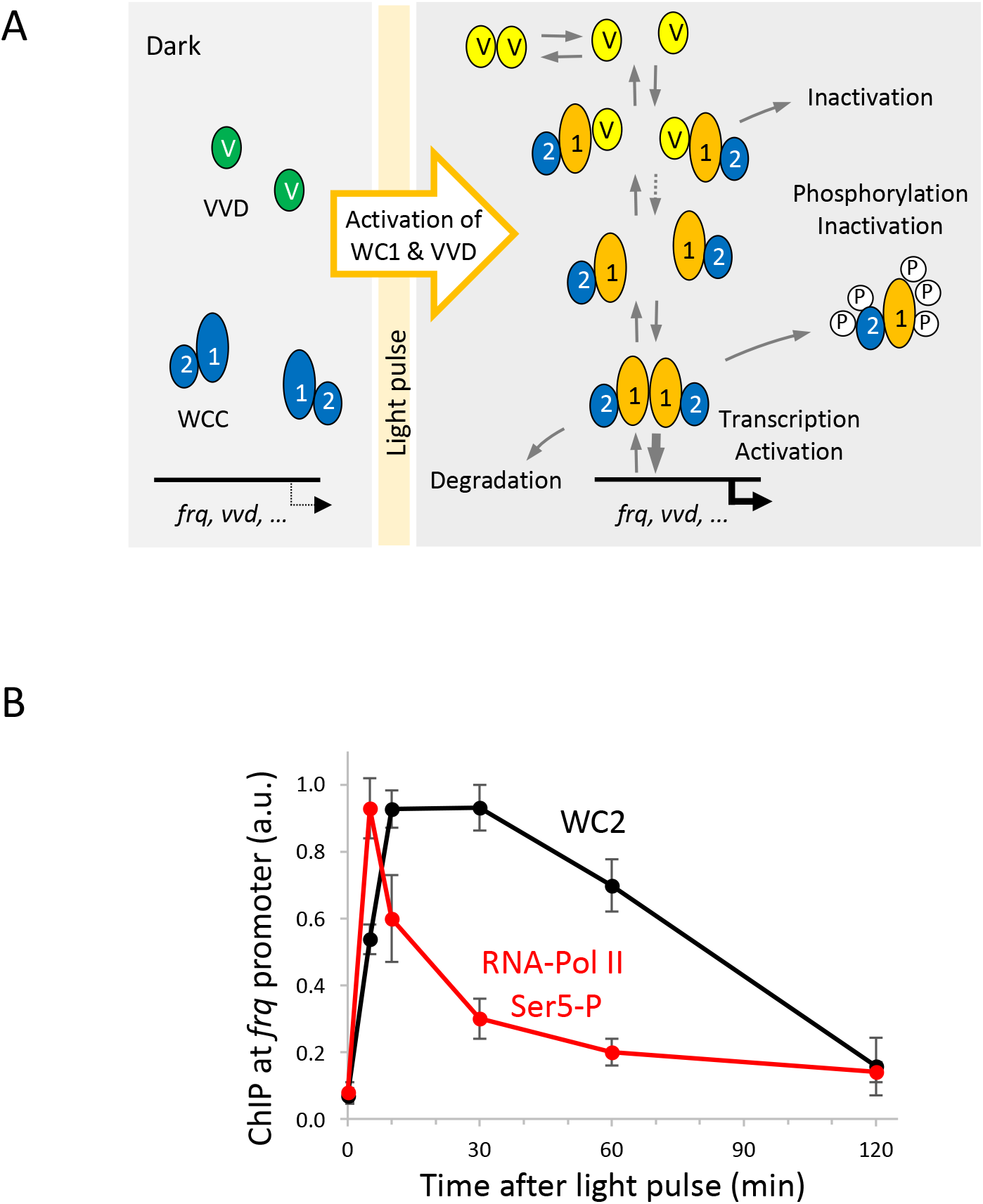
Promoter binding and induction of transcription by light-activated WCC. **A** Schematic representation of the transcriptional light response of *Neurospora crassa*. In the dark, the WCC is a heterodimeric protomer consisting of White Collar 1 (WC1) and WC2, and it supports low level rhythmic transcription of the core clock gene *frequency* (*frq*), *vivid* (*vvd*) and other clock-controlled genes. The dark form of WCC displays low affinity for so-called light-response elements (LREs), which are found in the control regions of light-inducible genes, including *frq* and *vvd* (*50-52*). Blue-light exposure induces in the Light-Oxygen-Voltage-(LOV)- domain of WC1 a stable (t_1/2_ ∼3 h) cysteinyl photo-adduct with its FAD cofactor and triggers a conformational change that allows the dynamic dimerization of two WCC protomers, which then bind to LREs and activate transcription (*28, 50, 53, 54*). The activity of the light-activated WCC is rapidly attenuated by light-induced phosphorylation (*55, 56*), by degradation (*57-59*), but mainly by interaction with VVD (*28-30*), a small protein consisting essentially of a single LOV domain (*31*). Light-activated VVD dynamically forms homodimers (*60*) and heterodimers with WCC via the light-activated LOV domain of WC1 (*28*). Thus, VVD competes with WCC homodimerization and efficiently attenuates light-dependent transcription (*28, 30*), although the mechanism is not fully understood. Phosphorylation of WCC and interaction with VVD are not mutually exclusive. **B** Transcription activated by a light pulse (LP) ceases rapidly despite persistent binding of WCC. Dark-grown *Neurospora* cultures were exposed to a 1-min high-LP. Transcriptional activity of the *frq* promoter and WCC binding to the *frq* LRE were measured by Pol II Ser5p ChIP-qPCR (data from (*21*)) and WC2 ChIP-qPCR, respectively, before and at the indicated time periods after the LP (mean values ±SEM, *n* = 3).

Here, we asked whether WCC impacts light-induced gene expression beyond its known function as transcription activator. We modulated WCC activity and abundance *in vivo* and analyzed the impact on transcription of luciferase reporter genes controlled by *frq* and *vvd* promoters. Our results suggest that WCC has also a role as a transcription repressor. Repression occurs in the wake of transcription activation and is facilitated by the interaction of WCC with histone deacetylase 3 (HDA3), the catalytic subunit of large co-repressor complexes such as RPD3(L/S) in yeast and SMRT/NCoR in mammals (*23-27*). Our data suggest that transcriptional bursts are temporally limited by co-repressors recruited by the WCC.

## RESULTS

### Discrepancy between WCC activity and transcription

Activation of the WCC by a light pulse (LP) recruits WCC to light-response elements (LREs) and triggers transcription from target promoters such as *frq* and *vvd* as schematically outlined in Fig. 1A. However, as shown for the *frq* promoter (Fig. 1B), the LP-induced Pol II transcription wave declines rapidly even though analysis by ChIP indicated that WCC occupies the *frq* LRE for a much longer period of time. The data suggest that the pool of initially activated WCC loses its ability to activate transcription before eventually becoming incompetent for DNA binding or being degraded. Interestingly, WCC occupancy of the *frq* promoter correlates fairly well with the refractory state of the *frq* promoter (t_1/2_ ∼45 min) that is established in the wake of transcription (*20, 21*). We wondered how the prolonged occupancy of LREs by light activated WCC, which is highly dynamic (*21*), was related to the observed decrease in transcription. Thus, we set out to manipulate the duration and extent of WCC activity *in vivo*. Levels of activated WCC decrease rapidly due to progressive inactivation by VVD (*28-30*), which ultimately terminates LP-induced transcription waves. Because light-activated WCC is active longer in the absence of VVD (*31, 32*), we generated *Δvvd* strains and corresponding *vvd*^*+*^ strains for control (*vvd*^*+*^*ctrl*) expressing destabilized luciferase (Luc^PEST^) from the *vvd* and the *frq* promoter, respectively (*33*). The stability of the covalent WC1 LOV photo-adduct makes it possible to manipulate by a short light pulse (LP) of suitable intensity the cellular concentration of activated WCC over more than two orders of magnitude and thus analyse transcription dynamics *in vivo* as a function of TF concentration (*20, 21*). The *vvd-luc*^*PEST*^ and *frq-luc*^*PEST*^ reporter strains were exposed to a 1-min LP and bioluminescence was recorded. The LP intensity was stepwise halved over a range from 85 to approximately 0.04 μmol photons m^-2^ s^-1^ to activate different amounts of the cellular WCC pool and, in the control strain, of VVD, too. The LP induced a wave of luciferase expression (Fig. 2A) that was delayed compared with Pol II transcription (compare Fig. 1B), reflecting translation and accumulation of the enzyme. At all LP intensities, the response of *vvd-luc*^*PEST*^ was higher in *Δvvd* and lasted much longer than in the corresponding *vvd*^*+*^ background, *vvd*^*+*^*ctrl* (Fig. 2A). Moreover, expression of *vvd-luc*^*PEST*^ in *vvd*^*+*^*ctrl* peaked in a narrow range between 75 - 90 min, while in *Δvvd* the expression peak of *vvd-luc*^*PEST*^ shifted with increasing LP-intensity from 90 up to 180 min, indicating that WCC is active for a prolonged time period in absence of VVD. At LPs of 42.5 and 85 μmol photons m^-2^ s^-1^, the expression of *vvd-luc*^*PEST*^ in *Δvvd* increased biphasically, with a steep increase in the first 30 min followed by a slightly flatter increase. The observation suggests that high levels of WCC may partially attenuate *vvd* promoter transcription.

**Fig. 2.**
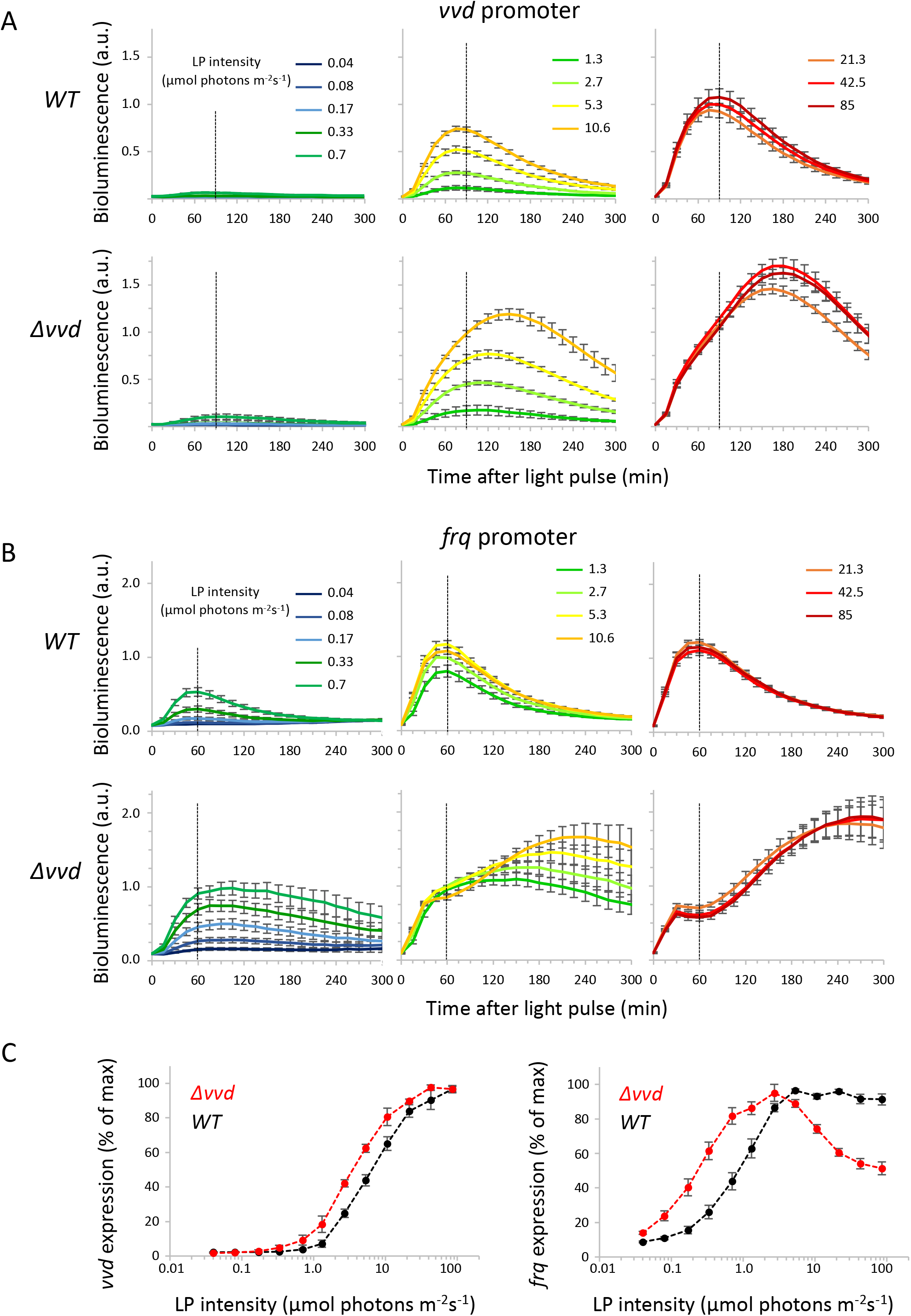
High levels of activated WCC attenuation *frq* transcription in *Δvvd*. Response of *vvd* and *frq* promoters to LPs of different intensity. Dark-grown (24 h) *WT* and *Δvvd* strains expressing **A**: *vvd-luc*^*PEST*^ and **B**: *frq-luc*^*PEST*^ were subjected to a 1-min LP of the indicated intensity. Luciferase activity was measured at 15-min intervals. Peak expression values of *frq-luc*^*PEST*^ at 2.7 μmol photons m^-2^ s^-1^ and of *vvd-luc*^*PEST*^ at 42.5 μmol photons m^-2^ s^-1^ (>80% saturation in *WT*) were normalized to 1 (mean values ±SEM, *n* = 4). Vertical dotted lines at 90 min (**A**) and 60 min (**B**), respectively, indicate approximately the expressions peaks in WT at saturating LP intensity. **C** Dependence of promoter activity on WCC concentration. The expression levels of *vvd-luc*^*PEST*^ after 90 min (left panel) and of *frq-luc*^*PEST*^ after 60 min (right panel) in *WT* (black curves) and *Δvvd* (red curves) were quantified from **A** and **B** and plotted against LP intensity. The highest value in each light intensity titration series was set to 100% (mean ±SEM, n = 4).

The expression level of *frq-luc*^*PEST*^ in the *vvd*^*+*^*ctrl* strain saturated at LP intensities >2.7 μmol photons m^-2^ s^-1^, and LP-induced expression waves peaked between 45 and 60 min at all LP intensities (Fig. 2B, upper panels). In *Δvvd* the LP-induced expression dynamics of *frq-luc*^*PEST*^ was surprisingly complex (Fig. 2B, lower panels). Low-intensity LPs (< 1.3 μmol photons m^-2^ s^-1^) induced expression waves that were higher, peaked later and lasted longer than in *vvd*^*+*^*ctrl*, consistent with higher and prolonged activity of WCC in absence of VVD (Fig. 2B, right panels). At intermediate LP intensities (1.3 - 10.6 μmol photons m^-2^ s^-1^) *frq-luc*^*PEST*^ expression increased biphasically, with a steep increase in the first 30 min (Fig. 2B, lower middle panel), similar to the transcription dynamics of *vvd-luc*^*PEST*^ at high LP-intensities (see Fig. 2B, lower right panel). However, at LP intensities ≥ 21.3 μmol photons m^-2^ s^-1^, *frq-luc*^*PEST*^ expression increased for the first 30 min but then did not reach the peak levels of *vvd*^*+*^*ctrl*. Rather, the expression decreased transiently and increased again after about 75 min to reach after 270 - 285 min peak levels above those of *vvd*^*+*^*ctrl* (Fig. 2B, lower right panel). Because the activity of WCC activated by a LP at 0 min decreases monotonically with time, the rebound of *frq-luc*^*PEST*^ expression after 75 min and the appearance of a second broad expression peak seems surprising at first glance. However, the observed two-peak transcriptional response to a single high-LP is consistent with transcriptional bursting of the *frq* promoter. The initial submaximal expression peak at 30 min suggests that supersaturation with high levels of WCC rapidly and synchronously represses the *frq* promoters in the population which then turn refractory to restimulation. Approximately 45 minutes later, when the repressed *frq* promoters begin to recover from the refractory state, there is still sufficient light-activated WCC present in *Δvvd* to trigger a detectable second and subsequently even further transcriptional bursts. However, with decreasing WCC levels, the second and further bursts of individual promoters are less synchronous due to stochastic TF binding, so that they partially overlap and are recognized as a broad wave of expression at the population level. Finally, when the activated WCC concentration falls below the saturation concentration the expression of *frq-luc*^*PEST*^ decreases. The time when levels of *frq-luc*^*PEST*^ begin decreasing in *Δvvd* correlate with the intensity of the LP administered at 0 min.

These data indicate that transcription from the *frq* promoter does not scale monotonically with the concentration of activated WCC. Rather, WCC has the potential to attenuate transcription (Fig. 2C), suggesting that WCC is not only an activator but additionally a transcription repressor. In our experimental setup, repression of the *frq* promoter is only visible at high concentrations of light-activated WCC because only then are LP-induced transcription, subsequent repression and recovery from the refractory state sufficiently synchronized to resolve them temporally at the population level. Furthermore, our data (Fig. 2A) suggest that WCC also represses the *vvd* promoter, but either less efficiently or for a shorter period than the *frq* promoter.

Transcription of *vvd* is activated by a LRE about 1.9 kb upstream of the core promoter (Fig. S1A), and we have previously shown that the potential to induce refractoriness at the *frq* promoter decreases when the LRE is moved in an upstream position (*20*). Analyzing previously published time-resolved Pol II ChIP-seq data (*21*), we noticed that the transcription of LP-induced noncoding RNAs directed directly from the *vvd* LRE was refractory to restimulation after 30 min, while *vvd* transcription, controlled by the same LRE was not (Fig. S1B). Hence, in addition to promoter architecture, proximity of the LRE to the core promoter and thus the local concentration of WCC appears to favour promoter refractoriness.

### WCC overexpression attenuates *frq* transcription

To further characterize the repressive role of WCC, we sought to increase the activity of light-activated WCC by overexpression but to limit the duration of its activity by VVD. Therefore, we generated *vvd*^*+*^ strains with a *frq-luc*^*PEST*^ reporter that overexpressed WC1 via the strong *ccg1* promoter (*wc1-overex*). The *wc1-overex* and a control strain expressing WC1 from the endogenous locus (*wc1-ctrl*) were exposed to a 1-min LP of increasing intensity and bioluminescence was recorded over a 300-min period (Fig. 3). As expected, expression waves in *wc1-ctrl* increased with LP intensity and saturated at ≥ 2.7 μmol photons m^-2^ s^-1^ (Fig. 3A, upper panels). Exposure of *wc1-overex* to LPs ≤ 2.7 μmol photons m^-2^ s^-1^ induced a higher *frq-luc*^*PEST*^ expression wave than in *wc1-ctrl* and reached a maximum at 1.3 - 5.3 μmol photons m^-2^ s^-1^ (Fig. 3A, lower panels). At higher light intensity, *wc1-overex* responded to a single LP with two successive expression peaks of *frq-luc*^*PEST*^, a first sharper peak at ∼30 min followed by a trough at ∼75 min and a second, broader peak with a maximum at 135 - 180 min (Fig. 3A, lower right panels). The temporal expression profile resembled the *frq-luc*^*PEST*^ dynamics in *Δvvd* (see Fig. 2B, C), except that the duration of the second expression wave was temporally more restricted in *wc1-overex* due to the presence of VVD. The observed two-peak transcriptional response to a single high-LP is consistent with transcriptional bursting of *frq* promoters.

**Fig. 3.**
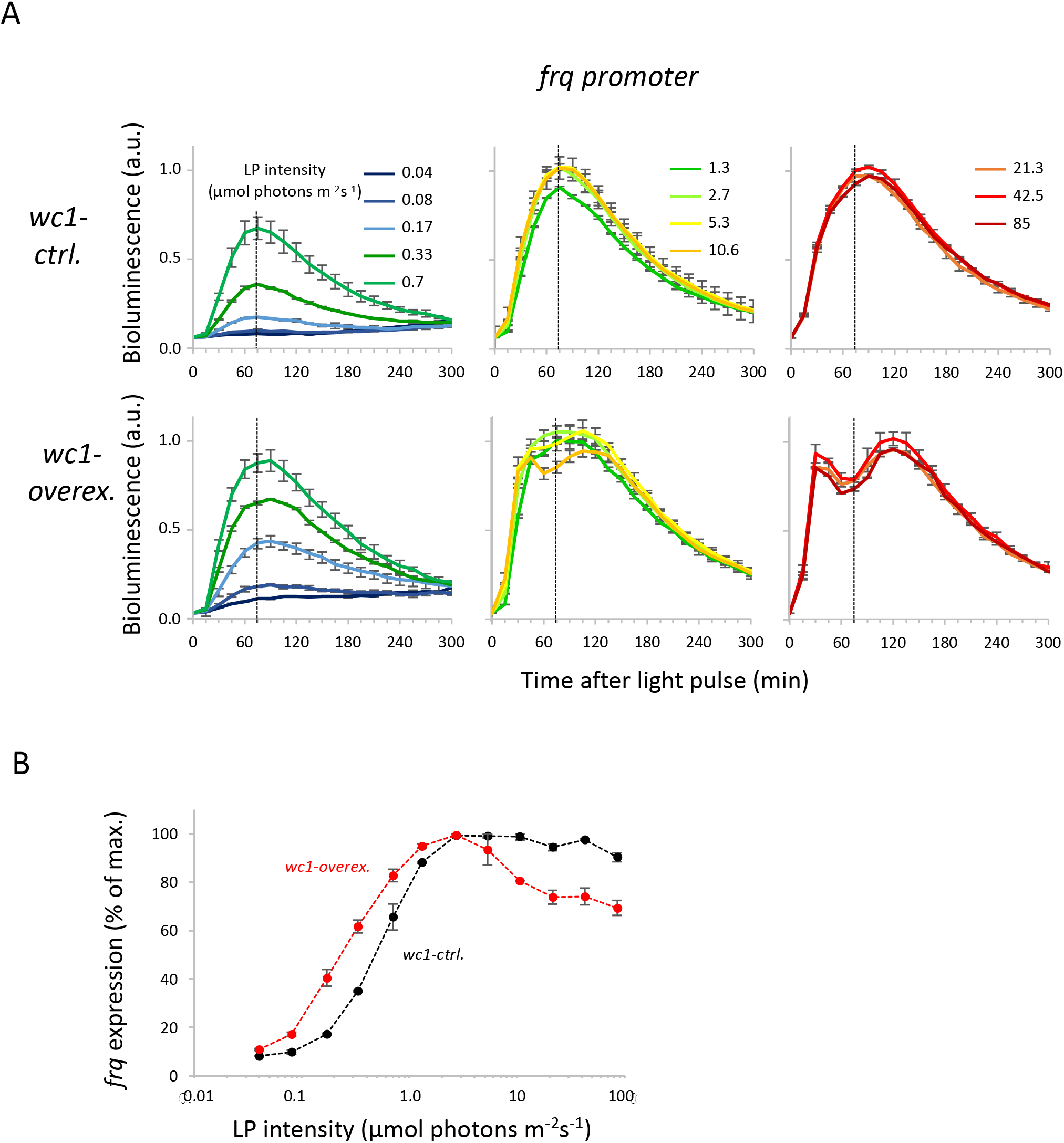
High WCC levels support two consecutive *frq* transcription waves in response to a single LP. **A** *WT* (*wc1-ctrl*) and strains overexpressing WC1 from the *ccg1* promoter (*wc1-overex)* were subjected to a 1-min LP of the indicated intensity. Luciferase activity was measured (mean values ±SEM, *n* = 2). Vertical dotted lines at 75 min indicate expression peaks at saturating LP intensities. **B** Dependence of promoter activity on WCC concentration. Expression levels of *frq-luc*^*PEST*^ 75 min after the LP were plotted versus LP intensity. Maximal values were set to 100 % (mean values ±SEM, *n* = 2).

### Repression of *frq* transcription correlates with WCC activity

We next sought to limit the duration of light-activated WCC activity independent of VVD by manipulating the stability of the photoadduct of the WC1-LOV domain. It was previously shown that two substitutions in the LOV domain of VVD, I_74_V together with I_85_V, result in a LOV photoreceptor with an unstable, rapidly decaying photoadduct (*34*). To this end, we generated a WC1 version with an I_405_V substitution in its LOV domain which corresponds to the I_85_V substitution in VVD. I_74_ of VVD corresponds to V_394_ of WC1 and was therefore not changed. WC1-I_405_V and a WT version of WC1 as a control were expressed in a *Δvvd* strain with a *frq-luc*^*PEST*^ reporter. The LP-induced expression waves of *frq-luc*^*PEST*^ in *Δvvd* strains expressing WC1-I_405_V were much shorter than in *Δvvd* strains expressing a WT version of WC1 (Fig. 4A and fig. S2), suggesting that the I_405_V substitution destabilized the WC1 photoadduct, as intended. Interestingly, expression dynamics of *frq-luc*^*PEST*^ induced by WC1-I_405_V in *Δvvd*, resembled that in *WT* strains, i.e. strains expressing WC1 and VVD (compare Fig. 4A right panel and Fig. 2C right panel). Thus, inhibition of WCC by VVD in WT appears to be similar to a reduction in WC1 photoadduct stability.

**Fig. 4.**
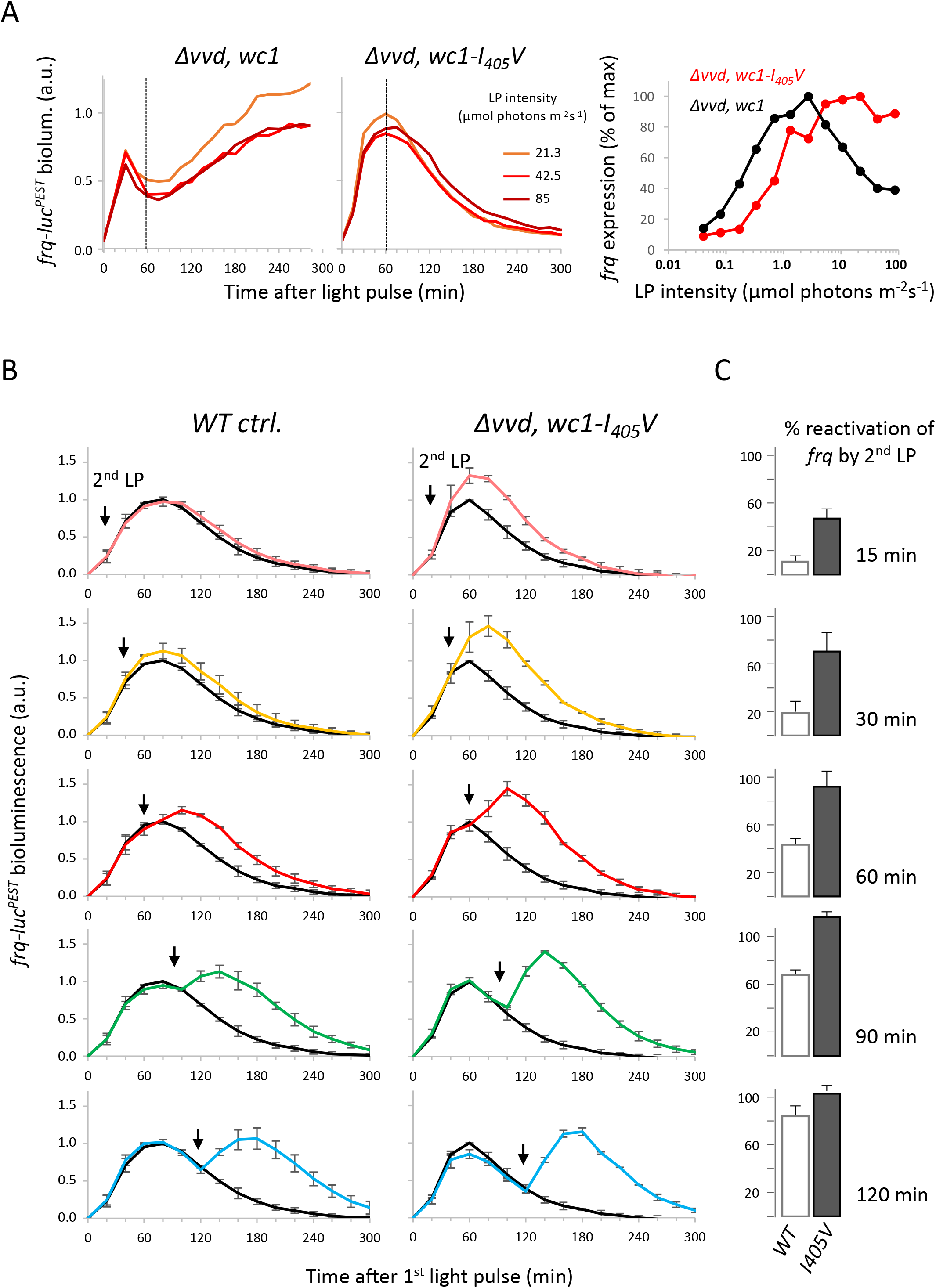
WC1 I_405_V supports restimulation of the *frq* promoter by consecutive LPs. **A** WC1 with an I_405_V substitution, which is predicted to destabilize the LOV photoadduct (*34*), was expressed in a *Δvvd* background. A control strain, *Δvvd wc1*^*+*^(left panel) and *Δvvd wc1-I*_*405*_*V* (middle panel) expressing *frq-luc*^*PEST*^ were subjected to a 1-min LP of the indicated intensity. (Expression profiles at lower LP intensities are shown in fig. S2A) Right panel: Expression levels of *frq-luc*^*PEST*^ 60 min after the LP were plotted versus LP intensity. Maximal expression values in *Δvvd wc1*^*+*^ and *Δvvd wc1-I*_*405*_*V*, respectively, were set to 100% (n=1). At high LP intensities expression of *frq-luc*^*PEST*^ was repressed in *Δvvd wc1*^*+*^ (black curve) but not in *Δvvd wc1-I*_*405*_*V* (red curve). **B** Restimulation of *frq-luc*^*PEST*^ by WC1 (left panels) and WC1-I_405_V (right panels). Black traces: Dark grown *vvd*^*+*^ *wc1*^*+*^ (*WT ctrl)* and *Δvvd wc1-I*_*405*_*V* strains expressing *frq-luc*^*PEST*^ were subjected to a 1-min saturating LP (5 μmol photons m^-2^ s^-1^) and luciferase activity was recorded. Expression dynamics of *frq-luc*^*PEST*^ were similar in both strains. Coloured traces: A second LP of 85 μmol photons m^-2^ s^-1^ was applied at the indicated time points after the first LP. Expression of *frq-luc*^*PEST*^ was refractory to early restimulation in *WT ctrl* but responsive to a second LP in *Δvvd wc1-I*_*405*_*V*. (mean values ±SEM, *n* = 2.) **C** Gain of *frq-luc*^*PEST*^ expression in response to the 2^nd^ LP relative to the 1^st^ LP (mean values ±SEM, *n* = 2).

We then asked whether WC1-I_405_V could still induce the refractory state of the *frq* promoter. Thus, *Δvvd, wc1-I*_*405*_*V* and *WT ctrl* strains (*vvd*^*+*^, *wc1*^*+*^) were first exposed to a 5.3 μmol photons m^-2^ s^-1^ LP. The single LP was sufficient to saturate expression of *frq-luc*^*PEST*^ and induced an expression wave of similar magnitude and duration in both strains (Fig. 4B, black traces). Subsequently, the strains were exposed to a second LP with an intensity of 85 μmol photons m^-2^ s^-1^, which was administered at different times after the first LP (Fig. 4B, coloured traces). As previously reported, the activated *frq* promoter turns refractory to restimulation and recovers from refractory state with a half-life of approximately 45 min (*20, 21*). Accordingly, expression of *frq-luc*^*PEST*^ in the *wc1* control strain barely responded to a second LP administered after 15 min, and the response increased gradually the later the second LP was administered after the first (Fig. 4C). In contrast, expression of *frq-luc*^*PEST*^ in *Δvvd* expressing WC1-I_405_V was more responsive to a second LP at all time points (Fig. 4C). The data indicate that WC1 and WC1-I_405_V induce transcription from the *frq* promoter with equal efficiency, whereas WC1-I_405_V is impaired in its ability to efficiently establish the refractory state at the *frq* promoter. Accordingly, transcription-induced repression of the *frq* promoter appears to depend on the sustained activity of the light-activated WCC.

### WCC interacts with the transcriptional co-repressor Histone Deacetylase 3

We hypothesized that WCC must interact with a transcriptional co-repressor in order to attenuate transcription of *frq-luc*^*PEST*^. Since histone deacetylases are common components of several co-repressor complexes (*35*) we analyzed knock-out strains of the four histone deacetylases genes (*hda1, 2, 3, and 4*) encoded by the *Neurospora* genome (*36*). The four knockout strains, *Δhda1, Δhda 2, Δhda 3*, and *Δhda 4*, and a *WT* for control were exposed to a single LP (85 μmol photons m^-2^ s^-1^) and *frq* RNA synthesized after 15 and 30 min was quantified by RT-qPCR (Fig. 5A). In WT, light induced *frq* RNA levels peaked 15 min after the LP and then decreased at 30 min. *Δhda1, Δhda 2*, and *Δhda 4* strains displayed similar dynamics of *frq* expression. However, in *Δhda3* the levels of *frq* RNA were higher after 15 min and kept increasing 30 min after the LP, suggesting that in *WT*, HDA3 may contribute to the downregulation of *frq* transcription after light induction.

**Fig. 5.**
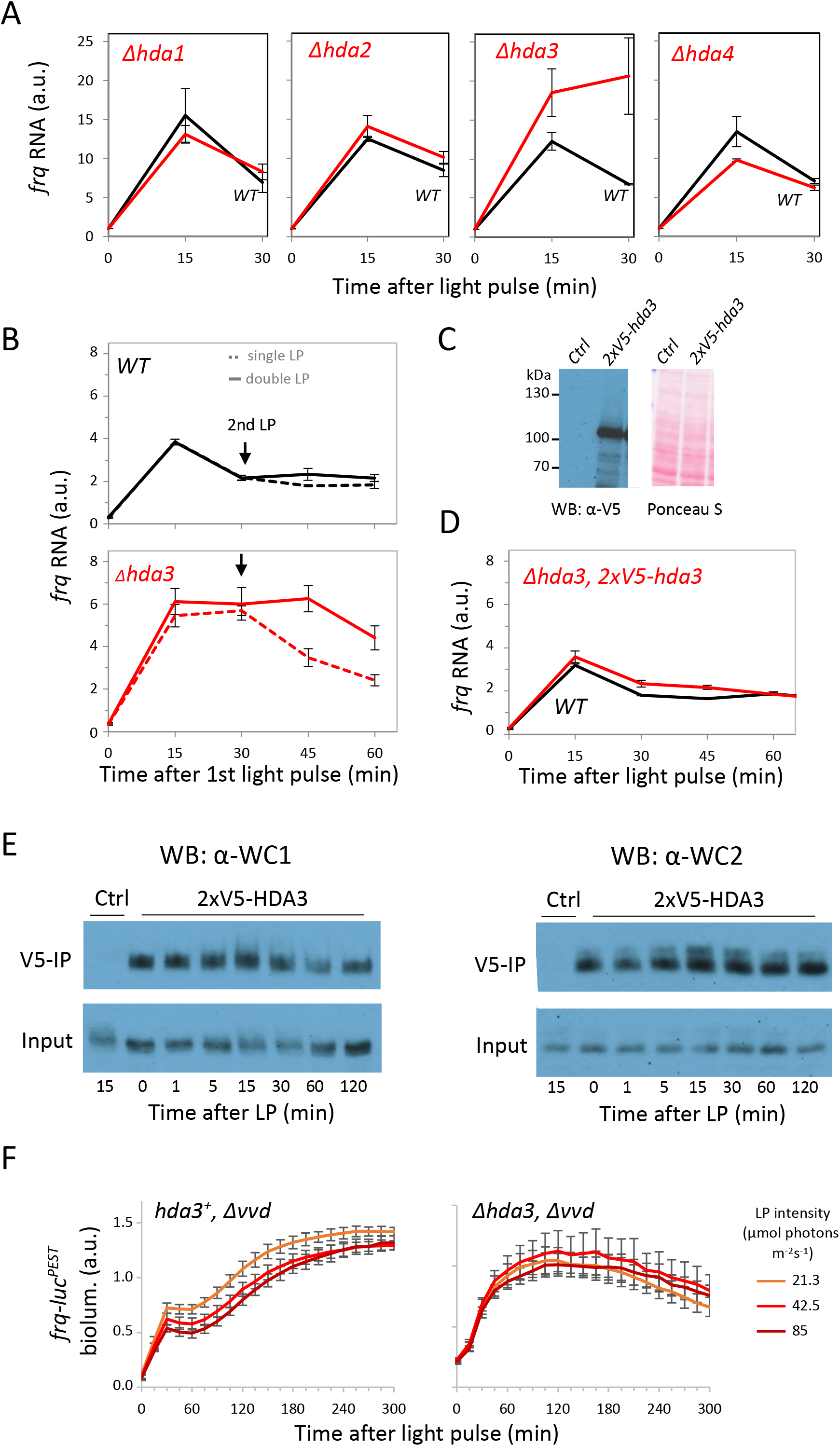
HDA3 interacts with WCC and attenuates light-induced transcription of *frq*. **A** Light-induced transcription of *frq* is elevated in a *hda3 KO* strain. Cultures of *WT* and *hda1, hda2, hda3* and *hda4* knockouts strains were subjected to a saturating 1-min LP. Samples were harvested at indicated time points. RNA was extracted and *frq* expression was measured by RT-qPCR. Expression levels were normalized to *actin* RNA. (mean values ±SEM, *n* ≥ 2). **B** Expression of *frq* is responsive to a 2^nd^ LP in *Δhda3* but refractory towards restimulation in *WT*. Cultures of *WT* and *Δhda3* were subjected at 0 min to a 1^st^ LP (5.3 μmol photons m^-2^ s^-1^) and at 30 min (arrow) to a 2^nd^ LP (85 μmol photons m^-2^ s^-1^). Samples were harvested at indicated time points and *frq* expression was measured. Data were normalized to 28S rRNA (±SEM, *n* = 3). **C** Overexpression of 2xV5-HDA3 under control of the *tcu1* promoter in *Δhda3*. Western blot analysis with V5 antibodies. Ctrl: *WT* strain. **D** Overexpression of 2xV5-HDA3 in *Δhda3* restores dynamics of *frq* induction. *WT* (Ctrl) and *Δhda3, ptcu1-2xV5-hda3* were grown in Cu^2+^-free medium to induce the *tcu1* promoter. Cultures were subjected to a 1-min LP (85 μmol photons m^-2^ s^-1^). Samples were harvested at the indicated time points and *frq* expression was measured by RT-qPCR. Expression levels were normalized to 28S rRNA (±SEM, *n* = 3). **E** HDA3 interacts with WCC in the dark and after light exposure. *Δhda3, ptcu1-2x-v5-hda3* and *WT* (*Ctrl*) were grown in Cu^2+^-free medium and exposed to a 1-min LP (85 μmol photons m^-2^s^-1^). Samples were harvested at the indicated time points. Native whole cell lysates were prepared and subjected to immunoprecipitation with V5 antibodies. WC1 and WC2 in input and IP samples were analyzed by Western blotting. WC1 and WC2 co-immunoprecipitated with 2x-V5-HDA3 at all time points (see also fig. S3A), but not from control extract (no tagged HDA3). **F** Absence of HDA3 prevents LP-induced transient repression of the *frq* promoter in *Δvvd. hda3*^*+*^*Δvvd* and *Δhda3Δvvd* strains expressing *frq-luc*^*PEST*^ were subjected to a 1-min LP of indicated intensity and luciferase activity was measured. (±SEM, *n* = 4). (Expression profiles at lower LP intensities are shown in fig. S3B)

Therefore, *Δhda3* and *WT* were treated with either a low LP (5.3 μmol photons m^-2^ s^-1^) or a low LP followed by a high LP (85 μmol photons m^-2^ s^-1^) after 30 min, and *frq* RNA was measured by RT-qPCR (Fig. 5B). The transcriptional activity of *frq* in response to a single LP was higher in *Δhda3* and lasted longer than in *WT*. As expected, *frq* transcription in *WT* barely responded to the second LP at 30 min, indicating that the *frq* promoter was repressed and refractory to restimulation (Fig. 5B, upper panel). In contrast, *frq* transcription in *Δhda3* responded to the second light pulse (Fig. 5B, lower panel). These data indicate that HDA3 contributes to the repression of transcription and the establishment of the refractory state at the *frq* promoter.

To determine whether HDA3 interacts with WCC, we expressed in *Δhda3* a version of HDA3 that was N-terminally tagged with a double V5 epitope (Fig. 5C). *Δhda3* exhibited a severe growth defect, and expression of 2xV5-HDA3 rescued the growth defect. Furthermore, levels and dynamics of LP-induced *frq* transcription were similar in *WT* and *Δhda3, 2xV5-hda3* (Fig. 5D). These data indicate that the tagged HDA3 protein was functional. We then prepared native protein extracts from dark-grown and light-pulsed mycelial cultures and performed pull-downs with V5 antibodies. Both WCC subunits, WC1 and WC2, co-precipitated with 2xV5-HDA3 in samples from dark-grown and LP-exposed samples (Fig. 5E and fig. S3A). In contrast, neither WC1 nor WC2 co-precipitated with V5 antibody from extract prepared from a *WT* control strain, expressing untagged HDA3. The data indicate that HDA3 is a constitutive interactor of WCC.

Finally, we generated a *Δhda3, Δvvd* double knockout strain containing a *frq-luc*^*PEST*^ reporter gene. This strain, and for comparison a *hda3*^*+*^, *Δvvd* strain with a *frq-luc*^*PEST*^ reporter gene, were exposed to LPs of different intensities (Fig. 5F and fig. S3B). Expression of *frq-luc*^*PEST*^ in response to a high-intensity LP was transiently repressed 30 - 60 min after exposure of *hda3*^*+*^, *Δvvd* (Fig.5F, left panel) in agreement with data shown in Fig. 2B. In contrast, the *Δhda3, Δvvd* strain showed no transient repression of *frq-luc*^*PEST*^ (Fig. 5F, right panel). The data indicate that HDA3 contributes to the downregulation of *frq* transcription upon activation of WCC. Finally, to assess whether HDA3 affects transcription of *vvd*, we exposed *WT* and *Δhda3* strains to a 1-min high LP and quantified *vvd* RNA before and at different times after the LP. The LP-induced levels of *vvd* RNA were higher in *Δhda3* than in *WT* (fig. S3C) indicating that HDA3 also attenuates *vvd* transcription.

## DISCUSSION

Here, we analyzed the transcription dynamics of the *frq* promoter and the approximately tenfold more potent *vvd* promoter, both of which are activated by the light-activatable transcription factor WCC of *Neurospora*. Upon activation by an LP, WCC dimerizes and rapidly binds to LREs located in the core promoter of *frq* and in an upstream activation sequence (UAS) of *vvd*, respectively. This triggers a synchronized wave of Pol II transcription in the cell population that peaks after approximately 15 minutes. However, while transcription stops rapidly, WCC keeps the LREs occupied for a much longer period, indicating that WCC no longer activates transcription. We show that the activating potential of WCC is rapidly antagonized by HDA3 in a promoter-specific manner. HDA3 is the *Neurospora* homolog of the conserved histone deacetylases Rpd3 and HDAC3, subunits of Rpd3(L/S) and NCoR/SMRT corepressor complexes in yeast and in vertebrates, respectively (*21, 23-27*). At the *frq* promoter, HDA3 facilitates the establishment of a long-lived repressed state called the refractory period, which lasts for about 45 minutes (*21*). We show by co-IP that HDA3 interacts with WCC in the dark and after an LP and that the refractory period correlates with the time that WCC occupies the *frq* LRE. In the absence of HDA3 *(Δhda3*), *frq* transcription continues as long as WCC occupies the LRE, indicating that WCC is in principle active during this time period. Also, during the refractory phase, newly activated WCC rapidly exchanges with previously activated WCC but does not induce *frq* transcription (*21*). In contrast, newly activated WCC readily reactivates transcription of *vvd*, suggesting that the inactive state of *vvd* is less resistant (refractory) to restimulation. We show that a substitution in the LOV domain of WC1, which is predicted to shorten the lifetime of LRE-binding-competent WCC, impairs the establishment of the refractory state of *frq*. These data suggest that WCC recruits HDA3 to repress the *frq* promoter and establish and maintain the refractory state as long as it occupies the LRE. However, WCC is a transcriptional activator. To induce transcription, light-activated WCC rapidly recruits NGF-1 (*37*), the *Neurospora* homolog of GCN5, the acetyltransferase subunit of the conserved SAGA coactivator. Therefore, WCC-dependent transcription appears to be determined by the balanced recruitment of coactivators and corepressors. A strong genetic interaction of Gcn5 and Rpd3 in yeast has been previously reported (*38*).

It remains unclear why WCC and HDA3 establish a refractory state in *frq*, whereas the inactive state of the *vvd* promoter is readily challenged by newly activated WCC. Differences in promoter architecture including the positioning and stability of relevant nucleosomes could account for these differences. The position of the LRE relative to the core promoter could also contribute to the promoter-specific differences. Since WCC displays similar affinity for the LRE in the *frq* core promoter and LRE in the *vvd* UAS, the effective WCC concentration at the core promoter might be a determinant for transcriptional refractoriness. We have recently shown that the distance of the LRE from the core *vvd* or *frq* promoter correlates with the concentration of activated WCC required for half-maximal activation (*20*). An LRE can directly bind only one dimeric WCC. However, it has been shown that low-complexity domains in TFs promote the formation of dynamic hubs composed of multiple TFs or even larger assemblies (*39-42*). Consistent with the formation of dynamic TF hubs, it has been shown in several cases that the residence times of TFs at specific binding sites are rather short and exhibit a fairly broad power-law distribution (*43*). The light-induced formation of WCC hubs at LREs could account for the differences in the transcriptional dynamics of *frq* and *vvd* promoters as depicted in the model shown in fig. S4. A dependence of hub size on the concentration of activated WCC would also explain why the apparent binding of WCC to LREs, as measured by random formaldehyde cross-linking and subsequent ChIP, increases with light intensity even beyond conditions where the *frq* promoter is functionally saturated (*21*).

We analyzed transcriptional waves in response to LPs to decipher the mechanisms controlling the transcriptional dynamics of *frq* and *vvd*. Because the HDA3-dependent repressed state of *frq* is long-lived, antagonism between activating NGF-1 and repressing HDA3 should promote transcriptional bursting in steady state, i.e. constant light. The life-time of the repressed state of *frq* would be the major determinant of bursting. Indeed, Rpd(L) has been shown to repress transcription in yeast by decreasing the frequency of transcriptional bursts (*27*). Because the *frq* promoter is already functionally saturated at low WCC concentrations, the stochasticity of WCC binding should not affect bursting under most light conditions. In contrast, although *vvd* also is repressed by HDA3 (fig. S3C), the repression is short-lived. Because of the localization of its LRE in a UAS and the dynamics of enhancer-promoter looping, the vvd promoter is not saturated even at high WCC concentrations (*20*). Therefore, transcriptional dynamics and possibly burst frequency of *vvd* are predominantly determined by the stochasticity of WCC binding. Although HDA3 seems to also repress the *vvd* promoter, the repressed state is short-lived and our population approach does not allow sufficiently tight synchronization and temporal resolution of transcription waves to assess the duration of the repressed state of *vvd* by a second light pulse. In general, the transcriptional dynamics of highly tunable promoters like *vvd* should be determined mainly by the dynamics of TF binding. Strong promoters analyzed so far at the single cell level seem to fall into this category (*8, 12*). Transcriptional dynamics of promoters with pronounced refractory features (*4*) may be determined by an antagonism of activation and long-lived repression. Such promoters could, like *frq*, respond rapidly to signals in an almost digital all-or-nothing manner (*20*). Activation and repression could be mediated by the same TF, as shown here for the WCC. In other cases, transcription dynamics are regulated by TF-families of activators and repressors binding to the same regulatory elements, such as MYCs and MXDs or Rev/ERBs and RORs (*44-49*) or by the combinatorial activity of unrelated regulators.

## MATERIALS AND METHODS

### Plasmids

*frq-lucPEST* and *vvd-lucPEST* were described previously (*33*) Plasmids *wc1-I*_*405*_*V, wc1-overex and 2xV5 hda3* were constructed using standard techniques. Plasmid sequences are available in Supplementary information.

### *Neurospora crassa* strains

| *Strain* | *Genotype* | *Relevant phenotype* | *Reference/ origin* |
| --- | --- | --- | --- |
| WT *frq-lucPEST* | wt, *ras-1<sup>bd</sup>*, *his3::pfrq-lucPest-ttrpC* | wt with *frq-lucPEST* reporter | Cesbron et. al. 2013 (33) |
| WT *vvd-lucPEST* | wt, *ras-1<sup>bd</sup>*, *his3::pvvd-lucPest-ttrpC* | wt with *vvd-lucPEST* reporter |  |
| Δ*vvd frq-lucPEST* | *ras-1<sup>bd</sup>*, Δ*vvd::HygB<sup>r</sup>*, *his3::pfrq-lucPest-ttrpC* | *vvd* knockout with *frq-lucPEST* reporter | |
| Δ*vvd vvd-lucPEST* | *ras-1<sup>bd</sup>*, Δ*vvd::HygB<sup>r</sup>*, *his3::pvvd-lucPest-ttrpC* | *vvd* knockout with *vvd-lucPEST* reporter | |
| *pccg1-wt-wc1 frq-lucPEST*<br>(*wc1-overex*) | *ras-1<sup>bd</sup>*, Δ*wc1::HygB<sup>r</sup>*, Δ*wc2::HygB<sup>r</sup>*, *his3::pccg1-wt-wc1-ttrpC_pwc2-wc2-twc2_pfrq-lucPEST-tvvd* | *wcc* knockout with exogenous WT-WC1, WT-WC2 and *frq-lucPEST* reporter expression | This study |
| Δ*vvd pccg1-wt-wc1-V5 frq-lucPEST* | *ras-1<sup>bd</sup>*, Δ*wc1::HygB<sup>r</sup>*, Δ*vvd::HygB<sup>r</sup>*, *his3::pccg1-wt-wc1-ttrpC_pfrq-lucPest-tvvd* | *wc1 vvd* double knockout with exogenous WT-WC1-V5 expression and *frq-lucPEST* reporter | This study |
| Δ*vvd pccg1-I405V-wc1-V5 frq-lucPEST* | *ras-1<sup>bd</sup>*, Δ*wc1::HygB<sup>r</sup>*, Δ*vvd::HygB<sup>r</sup>*, *his3::pccg1-I405V-wc1-ttrpC*, *tub2::pfrq-lucPest-ttrpC* | *wc1 vvd* double knockout with exogenous I405V-WC1-V5 expression and *frq-lucPEST* reporter | |
| *pccg1-wt-wc1-V5 frq-lucPEST* | *ras-1<sup>bd</sup>*, Δ*wc1::HygB<sup>r</sup>*, *his3::pccg1-wt-wc1-ttrpC*, *tub2::pfrq-lucPest-ttrpC* | *wc1* knockout with exogenous WT- | |
|  |  | WC1-V5 expression and vvd-luc reporter |  |
| WT *ras-1<sup>wt</sup>* | wt, *ras-1<sup>wt</sup>* | WT | (FGSC #2489) |
| $\Delta hda3$ FGSC | *ras-1<sup>wt</sup>*, $\Delta hda3::HygB^r$, heterokaryon | *hda3* knockout | (FGSC #11614) |
| $\Delta hda3$ | *ras-1<sup>wt</sup>*, $\Delta hda3::HygB^r$ | *hda3* knockout | This study ($\Delta hda3$ FGSC was crossed with WT *ras-1<sup>wt</sup>*) |
| $\Delta hda3$ *ptcu1-2x-v5-hda3* | *ras-1<sup>wt</sup>*, $\Delta hda3::HygB^r$, *his3::ptcu1-2xV5-hda3-ttrpC* | *hda3* knockout with exogenous copper-tuneable V5-HDA3 expression | This study ($\Delta hda3$ FGSC was crossed with WT *ras-1<sup>wt</sup>* *ptcu1-2x-v5-hda3*) |
| $\Delta vvd$ *pfrq-luc ras-1<sup>wt</sup>* | *ras-1<sup>wt</sup>*, $\Delta vvd::HygB^r$, *his3::pfrq-lucPest-ttrpC* | *vvd* knockout with frq-luc reporter | This study ($\Delta vvd$ *ras-1<sup>wt</sup>* *his<sup>-</sup>* was transformed with pFH62_frq-lucPEST and pFH62_vvd-lucPEST) |
| $\Delta hda3\Delta vvd$ *pfrq-lucPEST* | *ras-1<sup>wt</sup>*, $\Delta hda3::HygB^r$, $\Delta vvd::HygB^r$, *his3::pfrq-lucPest-ttrpC* | *hda3/vvd* double knockout with frq-luc reporter | This study ($\Delta hda3$ FGSC was crossed with $\Delta vvd$ *pfrq-luc ras-1<sup>wt</sup>*) |
| $\Delta hda1$ | *ras-1<sup>wt</sup>*, $\Delta hda1::HygB^r$ | *hda1* knockout | (FGSC #12003) |
| $\Delta hda2$ | *ras-1<sup>wt</sup>*, $\Delta hda2::HygB^r$ | *hda2* knockout | (FGSC #11158) |
| $\Delta hda4$ | *ras-1<sup>wt</sup>*, $\Delta hda4::HygB^r$ | *hda4* knockout | (FGSC #11175) |

### Primers and TaqMan probes (Sigma Aldrich) for qPCR

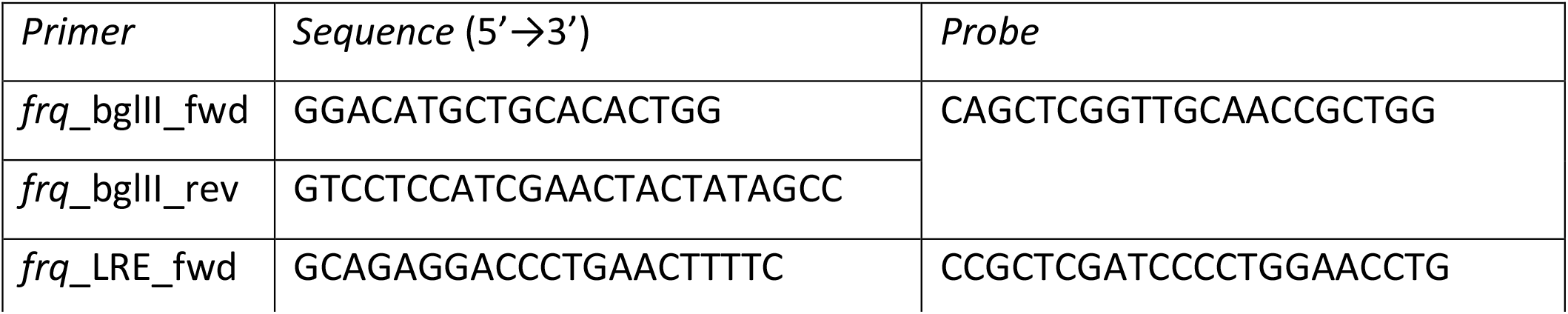
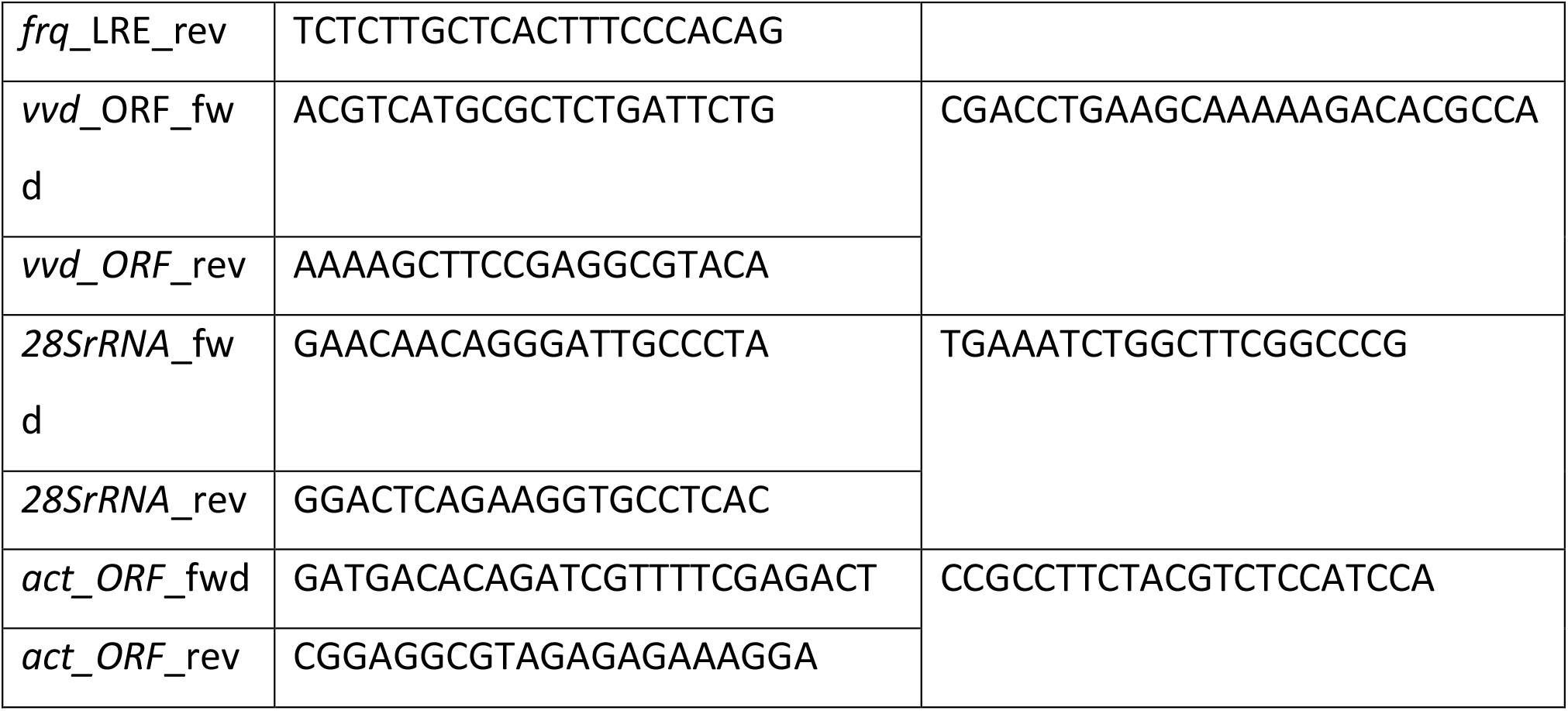

### *In vivo* luciferase assays

*Neurospora* was grown on solid medium (1x Vogel’s, 0.05 % [w/v] glucose, 0.05 % [w/v] fructose, 1.8 % [w/v] agarose and 10 ng/ml biotin, supplemented with 50 μM firefly luciferin) at 25°C. A 96-well plate containing 150 μl medium per well and was inoculated with 5μl of a 2.2 OD conidia suspension (ca. 30,000 conidia) and the plate was sealed with foil. Cultures were grown in constant light for 18 h, transferred to dark for 24 h and then exposed for 1-min to white light of indicated intensity. The light intensity was adjusted using light absorbing foil (E-colour+, #209, 3ND; Rosco, Stamford, USA). Light intensity is halved by each layer of light absorbing foil (*21*). Bioluminescence was recorded in darkness at 25 °C with an Perkin Elmer EnSpire Multilabel Reader. When indicated cultures were exposed to a second LP.

### Protein extraction and analysis

For the extraction of *Neurospora* native protein, tissue/cells were ground in the presence of liquid nitrogen using a precooled pestle and mortar. The powder was suspended in extraction buffer (50 mM Hepes/KOH [pH 7.4], 137 mM NaCl, 10% [vol/vol] glycerol, 5 mM EDTA containing 1 mM PMSF, 5 μg/mL leupeptin, and 5 μg/mL pepstatin A). Protein concentrations were determined by NanoDrop (PeqLab).

For Western blot analysis *Neurospora* whole cell lysate was analyzed by SDS/PAGE and transferred to a nitrocellulose membrane (GE Healthcare; 0.45 μm, 10 Å∼ 150 cm). The membrane stained with Ponceau S to control for gel loading prior to antibody decoration.

### Antibodies

Generation and affinity purification of WC1, WC2 and Pol II Ser5-P antibodies was described previously (*21*). V5 monoclonal antibodies were purchased from Thermo Fisher Scientific (R96025).

### Immunoprecipitation of V5-tagged proteins

Whole cell lysate (12 mg protein) was adjusted to a total volume of 500 μl with extraction buffer. V5-tagged protein was precipitated with 20 μl anti-V5 beads (Anti-V5 Agarose Affinity Gel, Cat.- No. A7345, Sigma-Aldrich) rotating on a wheel for 2 h at 4 °C. The beads were pelleted (100 x g, 3 min, 4 °C) and the supernatant was discarded. The beads were washed four times with extraction buffer and the bound protein was eluted with 50 μl 2 x Laemmli-Buffer (120 mM Tris/HCl pH 6.8, 4 % [w/v] SDS, 10 % [v/v] β-mercaptoethanol, 20 % [v/v] glycerol, 0.04 % [w/v] bromphenolblue) at 95 °C for 5 min.

### Chromatin immunoprecipitation (ChIP)

Frozen mycelia powder was resuspended in cold ChIP Lysis Buffer (50 mM HEPES/KOH pH 7.5, 150 mM NaCl, 1 mM EDTA, 1 % [v/v] Triton X-100, 0.1 % [w/v] SDS, 0.1 % [w/v] DOC, 5 μg/ml leupeptin, 5 μg/ml pepstatin A, 1 mM PMSF). The suspension was kept on ice for 30 min, repeatedly mixed by vortexing and finally sonicated using the Covaris S220X (Covaris) with the following settings: 180 peak power, 20 duty factor, 200 cycles/burst, 3 min, 4 °C (DNA fragment size ≤ 2000 bp). The sonicated sample was centrifuged (15000 x g, 30 min, 4 °C) and the supernatants were transferred into fresh DNA-low binding tubes (Sarstedt, Cat.-No. 72.706.700). The protein concentration was measured using Bradford assay (Roti®-Quant, Carl Roth, Cat.-No. K015). An equivalent of 3.85 mg of total protein was adjusted to a total volume of 1.1 ml with ChIP lysis buffer. The solution was pre-cleared with 20 μl BSA- and Salmon Sperm DNA-blocked Protein A Sepharose beads for 1 h at 4 °C on a rotation wheel. Beads were pelleted at 1000 x g at 4 °C. 1 ml of the supernatant was transferred into a fresh DNA-low binding tube, 25 μl were stored as input sample at -80 °C. α-WC2 antibody (Pineda, Berlin) was added and the lysates were incubated over night at 4 °C on a rotation wheel. The next morning 20 μl BSA- and Salmon Sperm DNA-blocked Protein A Sepharose beads were added and the α-WC2 antibody complexes were precipitated for 2 h at 4 °C on a rotation wheel.

The beads were washed on ice twice with ChIP-Lysis Buffer, once with Lindet Buffer (10 mM TRIS/HCl pH 8.0, 0.25 mM LiCl, 1 mM EDTA, 1 % [v/v] NP-40, 1 % [w/v] DOC, 5 μg/ml leupeptin, 5 μg/ml pepstatin A, 1 mM PMSF) and three times with TE-Buffer (10 mM TRIS/HCl pH 7.5, 1 mM EDTA, 5 μg/ml leupeptin, 5 μg/ml pepstatin A, 1 mM PMSF).

The protein complexes were eluted from the beads with 300 μl ChIP Elution Buffer (0.5 % [w/v] SDS, 0.1 M NaHCO_3_) shaking in a Thermomixer with 1000 rpm at 65 °C for 15 min. The input sample (see above) was mixed with 275 μl ChIP Elution buffer and was treated accordingly.

The beads were pelleted and 380 μl of the supernatant was transferred into fresh DNA low-binding tubes. 10 μg DNase-free RNase A and NaCl (final concentration: 0.2 M) was added and the sample was incubated at 65 °C over night to reverse the crosslinking. To degrade residual protein 20 μg Proteinase K, 10 mM EDTA and 40 mM TRIS pH 8.0 were added the next day. The mixture was incubated for 1 h at 50 °C.

The DNA was extracted using the Promega PCR Cleaning Kit (Wizard® SV Gel and PCR Clean-Up System, Promega Cat.-No. A9285) as indicated in the manual. DNA was eluted from the columns with 120 μl (ChIP-samples) or 50 μl (input samples) 60 °C warm DNase-free water. DNA solutions were stored at -20°C.

### RNA extraction

Frozen mycelia powder was resuspended in peqGOLD TriFast mix (peqlab, Cat.-No. 30-2010) and RNA was extracted following the manufacturer’s instructions. RNA pellets were dissolved in an appropriate volume of RNase free water (supplemented with 0.4 U/μl RNase inhibitor [RiboLock, Thermo Scientific, Cat.-No. EO0384]). RNA concentration was measured using NanoDrop (PeqLab).

### cDNA synthesis

Total cDNA was synthesized in a reaction volume of 10 μl from 0.5 μg RNA using the Maxima First Strand cDNA Synthesis Kit (Thermo Scientific, Cat.-No. K1642) following the manufacturer’s instructions. cDNA was diluted 1:15 in nuclease-free water.

### Quantitative Real-Time PCR

Quantitative real-time PCR (qPCR) was performed using qPCRBIO Probe Mix Hi-ROX (Nippon Genetics, Cat.-No. PB20.22-51) in 96-well plates with the StepOnePlus real-time PCR system (Applied Biosystems) under thermocycling conditions as described below.

### qPCR Cycling Conditions

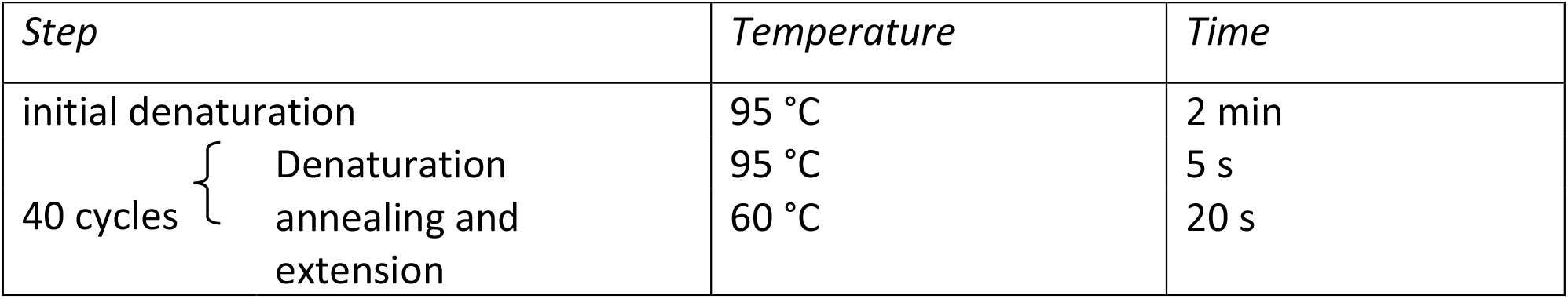

The reaction setup is described below. Mean cycle threshold (Ct) values were calculated from duplicates.

### qPCR Reaction Setup

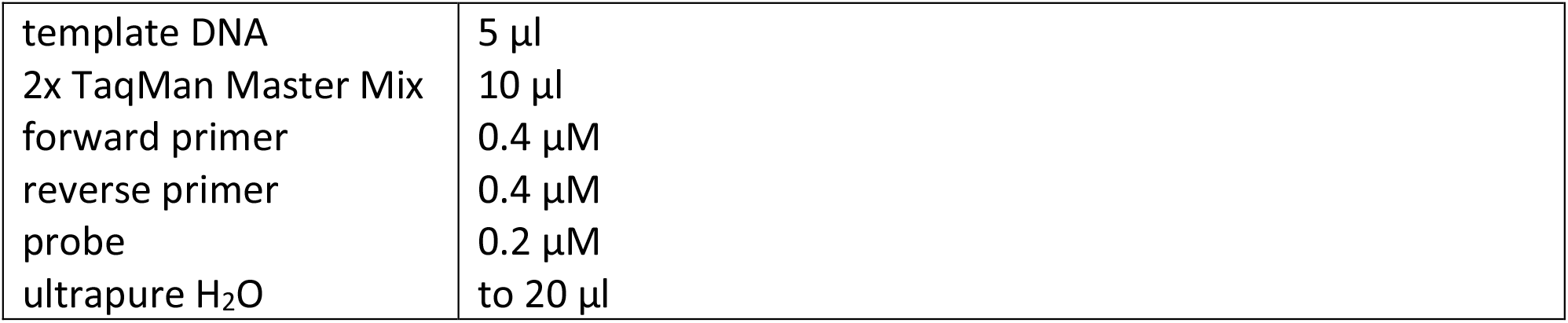

## ACKNOWLEDGEMENTS

We would like to thank the fungal genetics stock center (FGSC, Kansas, USA) for various Neurospora strains.

## Author contribution

Conceptualization: MB, MO; Methodology: MO, ACRD; Investigation: MO; Supervision: MB; Writing: MB, MO, ACRD

## Data and materials availability

ChIP seq data can be accessed via https://neutra.bzh.uni-heidelberg.de. All other data are present in the paper and/or the Supplementary Information.

## FIGURE LEGENDS

**fig. S1.**
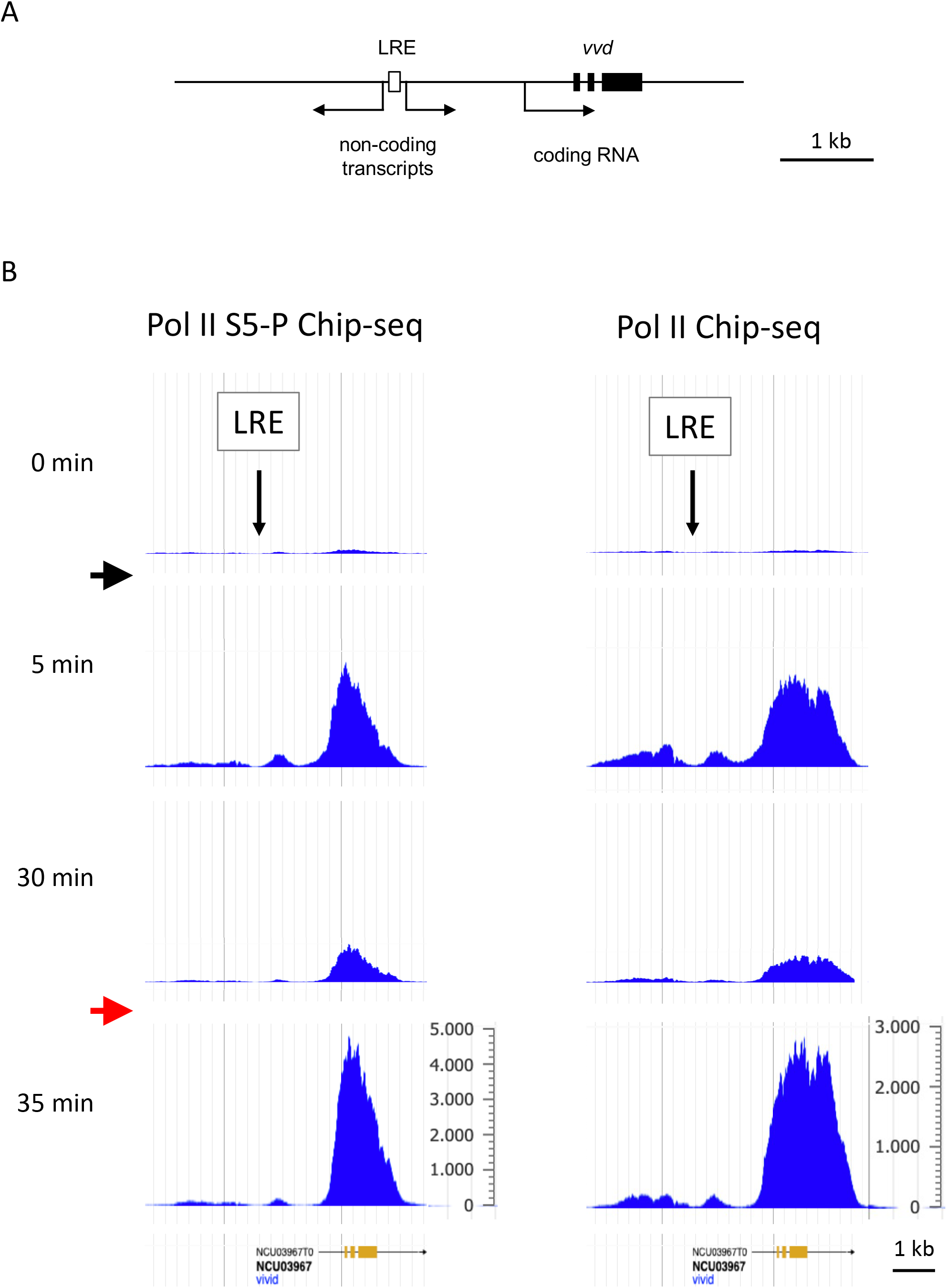
Differential response of transcription from *vvd* promoter and LRE. **A** Schematic of the *vvd* locus. The *vvd* LRE is located about 1.9 kb upstream of the promoter of the *vvd* coding gene. Non-coding transcripts originating at the *vvd* LRE are indicated. **B** Light-induced transcription originating directly from the *vvd* LRE is refractory to restimulation while transcription of the *vvd* coding gene, which is controlled by the same LRE remains responsive to restimulation. Dark-grown WT *Neurospora* was exposed to a 1-min low-LP (2.7 μmol photons m^-2^ s^-1^, black arrows) and after 30 min challenged by a 1-min high-LP (85 μmol photons m^-2^ s^-1^, red arrows). Samples were harvested at the indicated time points and subjected to ChIP-seq using antibodies against Pol II S5P (left part) and total Pol II (right part), respectively. Sequencing data are from (*21*) and accessible at https://neutra.bzh.uni-heidelberg.de.

**fig. S2.**
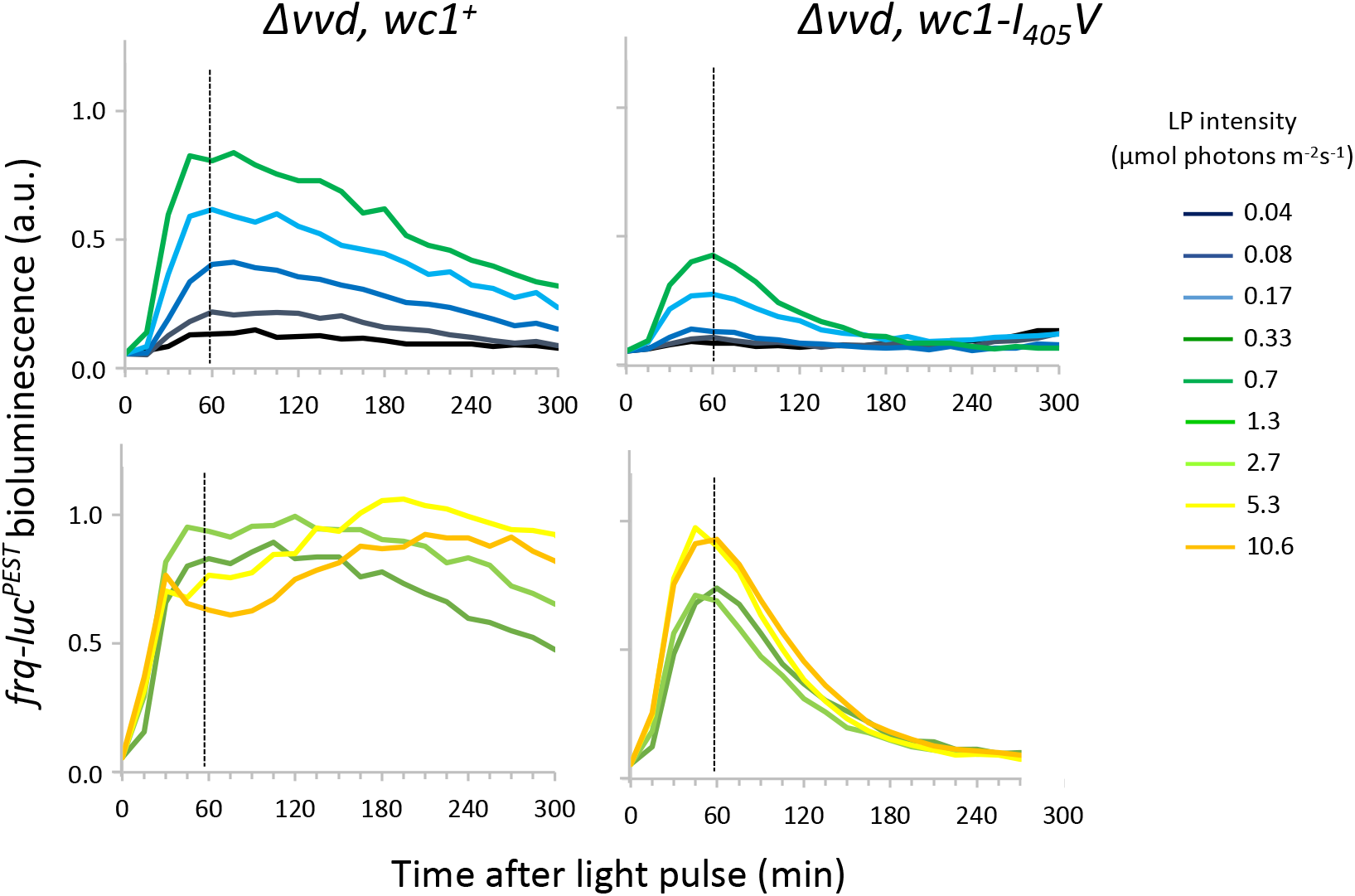
*frq-luc*^*PEST*^ expression dynamics in *Δvvd, wc1*^*+*^ (left panels) and *Δvvd wc1-I*_*405*_*V* (right panels)

**fig. S3.**
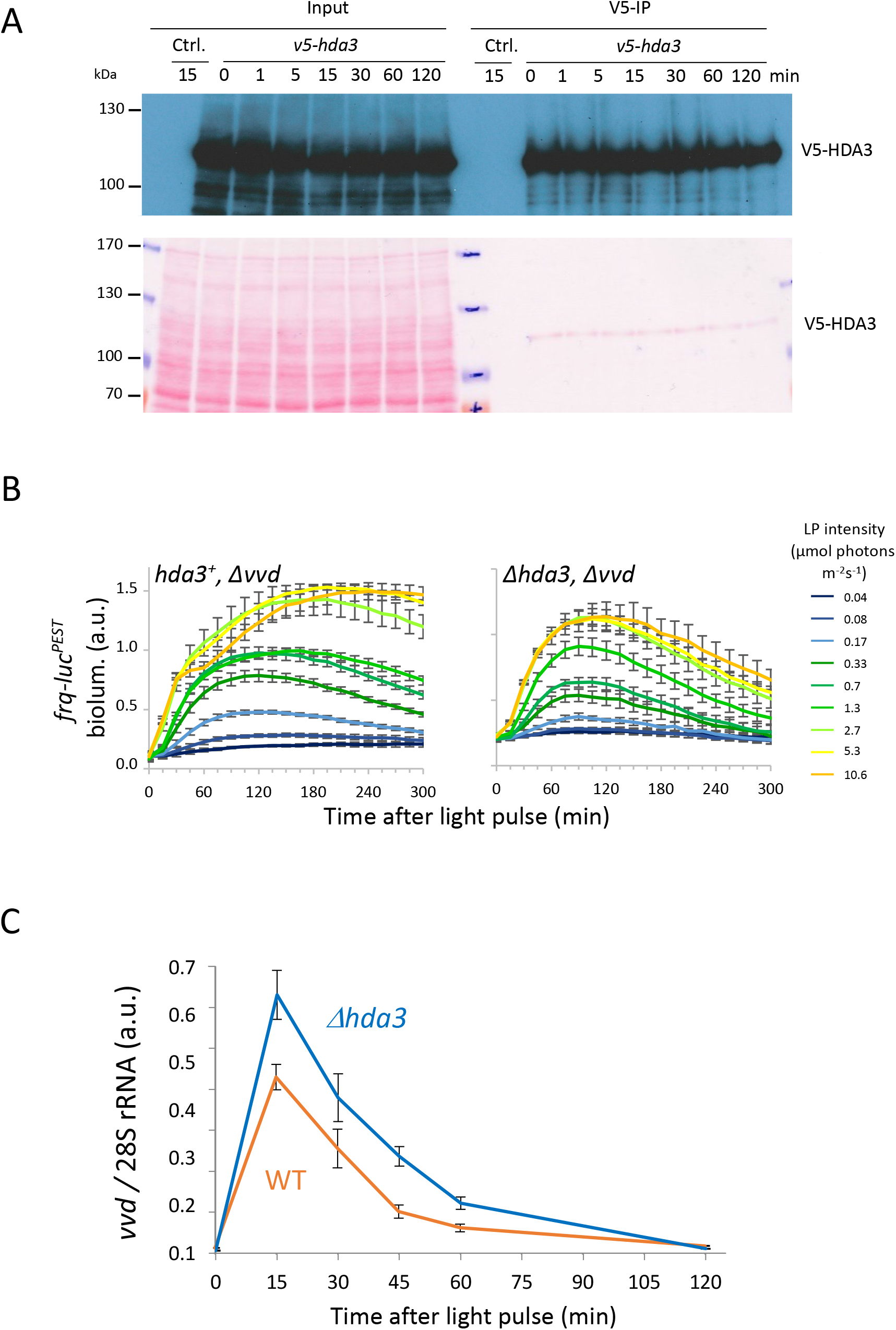
**A** V5 immunoprecipitation from WT and 2xV5-HDA3 overexpressing strains exposed to a LP at 0 min and harvested at the indicated time points (Fig. 5E). Upper panel: Samples were analyzed by Western Blot with V5 antibodies. Lower panel: Ponceau S stained membrane. **B** *hda3*^*+*^*Δvvd* and *Δhda3Δvvd* strains expressing *frq-luc*^*PEST*^ were subjected to a 1-min LP of indicated intensity and luciferase activity was measured. (±SEM, *n* = 4). Response of *frq-luc*^*PEST*^ to higher LP intensities is shown in Fig. 5F. **C** Expression of *vvd-luc*^*PEST*^ is elevated in absence of HDA3. WT and *Δhda3* were exposed to a 1-min LP (85 μmol photons m^-2^ s^-1^). Samples were harvested at the indicated time points and *vvd* expression was measured by RT-qPCR. Expression levels were normalized to 28S rRNA (±SEM, *n* = 3).

**fig. S4.**
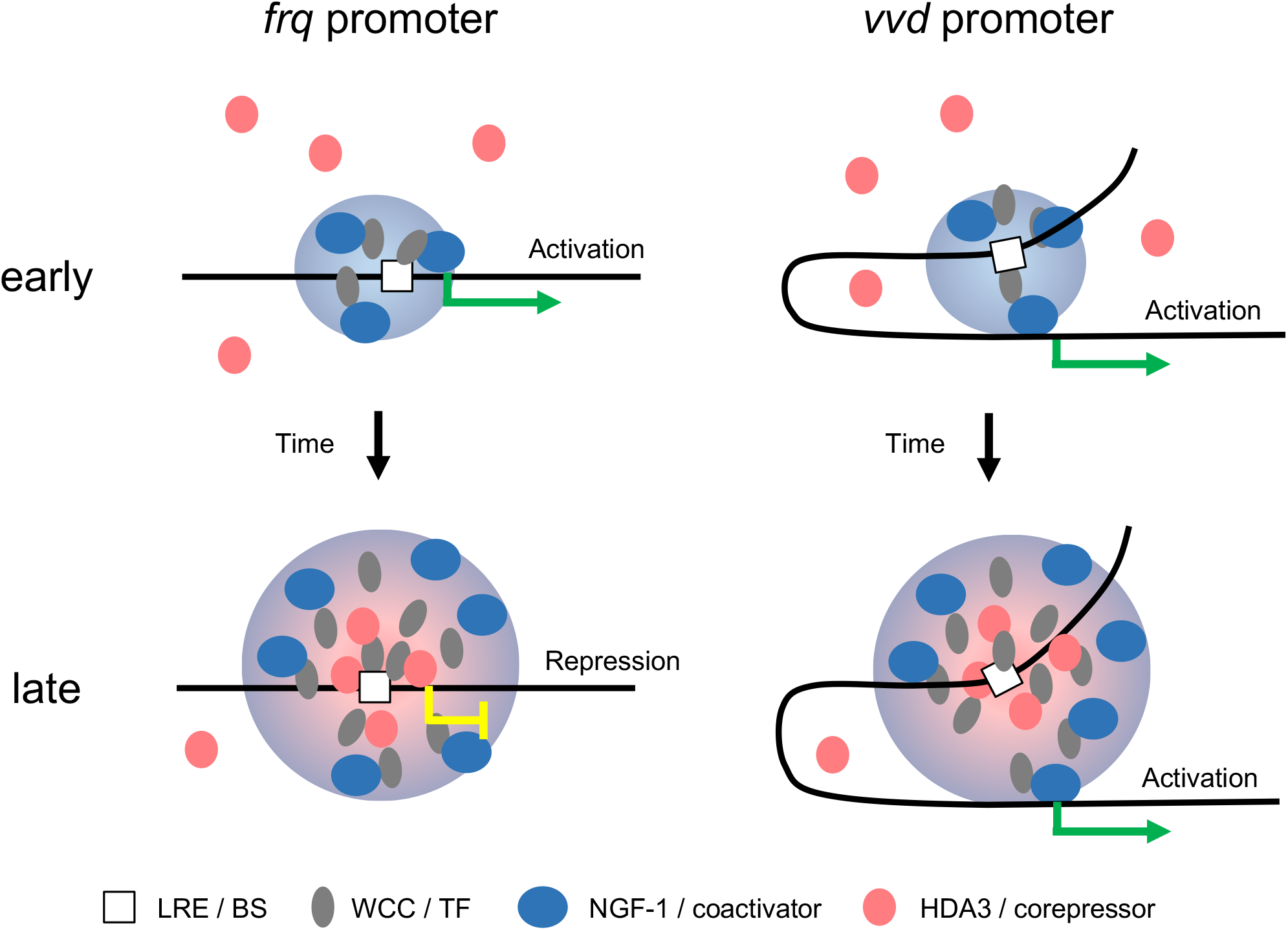
Molecular model for transcriptional refractoriness of the *frq* promoter to restimulation. The *frq* gene contains a LRE in the core promoter while the *vvd* LRE is located about 1.9 kb upstream of the promoter as schematically outlined. After light activation, WCC binds to the LREs and WCC hubs grow on the timescale of a transcription burst around an LRE that serves as a nucleation center. Due to the localization of their respective LRE, the *frq* promoter (left) is localized in the center of a WCC hub, whereas the *vvd* promoter dynamically interacts also with the hub surface through promoter-UAS looping. Provided HDA3 corepressor complexes have lower affinity for WCC than NGF-1 coactivator complexes, small WCC hubs do not efficiently sequester HDA3 and hence are activating. As WCC hubs grow over time, the interior of the hub provides more interaction sites for HDA3 and hence a more repressive environment than its surface, from which low-affinity interactors are easily lost. As it takes time to form WCC hubs of sufficient size to provide a repressive interior, the model explains why *frq* activation precedes repression. Additional WCC activated by a second LP (not depicted) would further increase the size of the dynamic hub, but the repressive environment inside would remain or even be enhanced, consistent with *frq* being refractory to restimulation. On the other hand, increasing hub size by a second LP would enlarge the activating hub surface around the *vvd* LRE and enhance transcription from the *vvd* promoter by increasing its probability to loop to its distantly located LRE. The observation that transcription originating directly from the *vvd* LRE is refractory to restimulation (fig. S1), whereas transcription from the more distant *vvd* promoter is not, supports this notion.

